# Marker-assisted pyramiding of two major broad-spectrum bacterial blight resistance genes, *Xa21* and *Xa33* into an elite maintainer line of rice, DRR17B

**DOI:** 10.1101/368712

**Authors:** CH Balachiranjeevi, S Bhaskar Naik, V Abhilash Kumar, G Harika, H.K Mahadev Swamy, Sk Hajira, T Dilip Kumar, M Anila, R.R Kale, A Yugender, K Pranathi, M.B.V.N Koushik, K Suneetha, V.P Bhadana, A.S Hariprasad, M.S Prasad, G.S Laha, G Rekha, S.M Balachandran, M.S Madhav, P Senguttuvel, A.R Fiyaz, B.C Viraktamath, A Giri, B.P.M Swamy, J Ali, R.M Sundaram

**Affiliations:** International Rice Research Institute, DAPO BOX 7777, Metro Manila 1301, Philippines; Biotechnology, Crop Improvement, Indian Institute of Rice Research, Indian Council of Agriculture Research, Hyderabad 30, India; Department of Plant Breeding, International Rice Research Institute-South Asia Hub, ICRISAT, Patancheru, India; Division of Crop Improvement, ICAR-Sugarcane Breeding Institute, Coimbatore, Tamil Nadu, India; Centre for Biotechnology, Institute of PG Studies and Research, Jawaharlal Nehru Technological University, Mahaveer Marg, Hyderabad 500 028, India.; ICAR-Indian Institute of Agricultural Biotechnology, PDU Campus, IINRG, Namkum, Ranchi, Jharkhand, India.

**Keywords:** Marker Assisted Backcross Breeding, Bacterial Blight, *Xa21* and *Xa33*, Hybrid rice, Maintainer line improvement, Host plant resistance

## Abstract

Bacterial blight (BB) disease reduces the yield of rice varieties and hybrids considerably in many tropical rice growing countries like India. The present study highlights the development of durable BB resistance into the background of an elite maintainer of rice, DRR17B, by incorporating two major dominant genes, *Xa21* and *Xa33* through marker-assisted backcross breeding (MABB). Through two sets of backcrosses, the two BB resistance genes were transferred separately to DRR17B. In this process, at each stage of backcrossing, foreground selection was carried out for the target resistance genes and for non-fertility restorer alleles concerning the major fertility restorer genes *Rf3* and *Rf4*, using gene-specific PCR-based markers, while background selection was done using a set of 61 and 64 parental polymorphic SSR markers respectively. Backcross derived lines possessing either *Xa21* or *Xa33* along with maximum genome recovery of DRR17B were identified at BC_3_F_1_ generation and selfed to develop BC_3_F_2_s. Plants harboring *Xa21* or *Xa33* in homozygous condition were identified among BC_3_F_2_s and were intercrossed with each other to combine both the genes. The intercross F_1_ plants (ICF_1_) were selfed and the intercross F_2_ (ICF_2_) plants possessing both *Xa21* and *Xa33* in homozygous condition were identified with the help of markers. They were then advanced further by selfing until ICF_4_ generation. Selected ICF_4_ lines were evaluated for their resistance against BB with eight virulent isolates and for key agro-morphological traits. Six promising two-gene pyramiding lines of DRR17B with high level of BB resistance and agro-morphological attributes similar or superior to DRR17B with complete maintenance ability have been identified. These lines with elevated level of durable resistance may be handy tool for BB resistance breeding.

## Introduction

Exploitation of heterosis for grain yield through hybrid rice technology is one of the feasible options to enhance rice production and productivity in order to meet the demands of an ever-increasing human population globally [1]. Even though rice hybrids were introduced in India in the early 1990s, their adoption has been slow and presently hybrid rice is cultivated in a limited area of 2.5 million ha. One of the principal reasons for slow adoption of rice hybrids in India is their susceptibility to major rice diseases like bacterial blight (BB) and blast [2]. Most of the commercial rice hybrids that have been released and cultivated in India do not possess broad spectrum resistance for BB disease [3].

BB disease is caused by a gram-negative bacterium called *Xanthomonas oryzae* pv. *oryzae* (*Xoo*). It is one of the most devastating diseases in rice [4]. The bacterium infects rice at maximum tillering stage, leading to water soaking lesions (blighting) on the leaves, which gradually enlarge, wilts and causes yield losses ranging from 74 to 81% [5]. Application of chemicals or antibiotics against BB is not effective [6, 7]. Breeding BB resistant rice varieties and hybrids is the best strategy for managing the BB disease in rice [8]. To date, at least 41 BB resistance genes have been identified, and some of them *viz*., *Xa4, xa5, xa13, Xa21* have been extensively used for development of BB resistant rice varieties [9, 10, 11]. However, large scale and long-term cultivation of varieties and hybrids with a single gene results in the breakdown of resistance due to a high degree of pathogenic variation [10, 12, 13]. Pyramiding of two or three *Xa* genes can enhance the durability and spectrum of resistance against BB [14, 15].

The major BB resistance gene, ‘*Xa21*’ was identified from *Oryza longistaminata*. It is located on chromosome 11 and a tightly linked to gene-specific marker pTA248 [16]. Similarly, ‘*Xa33*’ was identified from *Oryza nivara*. It is located on chromosome 7 and tightly linked to a marker RMWR7.6 [17]. These markers can be used in marker-assisted breeding to introgress *Xa21* and *Xa33* genes into different rice varieties and hybrid parental lines. These two genes are found to be highly effective against several isolates of *Xoo* from India and hence, are ideal choices for pyramiding into popular rice varieties or hybrids through marker-assisted breeding.

DRR17B is a promising maintainer line developed by ICAR-Indian Institute of Rice Research, Hyderabad, India [18]. It is however highly susceptible to BB of rice. In the present study, two major dominant BB resistance genes, *Xa21* and *Xa33* were introgressed into the genetic background of DRR17B through marker-assisted backcross breeding to develop improved DRR17B lines with broad spectrum resistance against BB.

## Materials and methods

### Plant materials

Improved Samba Mahsuri (ISM) is a recently released high-yielding and fine grain rice variety possessing BB genes, *xa5, xa13, and Xa21* [14]. It was used as a donor for *Xa21* [18]. A Near Isogenic Line (NIL) of Samba Mahsuri (FBR1-15EM) served as the donor for *Xa33* [17]. The popular but BB susceptible maintainer line DRR17B was used as a recurrent parent. It was developed by ICAR-Indian Institute of Rice Research (IIRR), Hyderabad, India.

### Strategy for marker-assisted introgression of *Xa21* and *Xa33* into DRR17B

Marker-assisted backcross breeding strategy was adapted for targeted introgression of *Xa21* and *Xa33* genes into the genetic background of the elite maintainer line of rice, DRR17B. Each of these genes was separately introgressed into DRR17B through two sets of crosses, i.e., Cross I, *viz*., DRR17B/Improved Samba Mahsuri and Cross II, *viz*., DRR17B/FBR1-15 (Fig 1). The F_1_s obtained from the two crosses were analysed using gene-specific markers pTA248 (specific for *Xa21*; [16]) and RMWR7.6 (specific for *Xa33*; [17]) to identify ‘true’ heterozygotes. The ‘true’ F_1_s were backcrossed with the recurrent parent DRR17B to generate BC_1_F_1_s, which were then screened for the presence of the target resistance genes using the gene-specific markers. The positive plants for *Xa21* and *Xa33* were selected and further screened for the non-presence of major fertility restorer genes, *Rf4* and *Rf3* using tightly linked markers, viz., DRCG-*RF4*-14 and DRRM-*RF3*-10, respectively [19]. BC_1_F_1_ plants possessing BB genes and a non-restoring allele concerning *Rf4* and *Rf3* in homozygous condition were selected following the procedure described by [18]. These plants were later screened with a set of parental polymorphic SSR markers (61 markers specific to the cross DRR17B/ISM and 64 markers specific for the cross DRR17B/FBR1-15EM) through background selection to identify a single BC_1_F_1_ plant from each cross possessing maximum recovery of the recurrent parent genome. The selected plant was backcrossed once again with DRR17B.

**Fig. 1.**
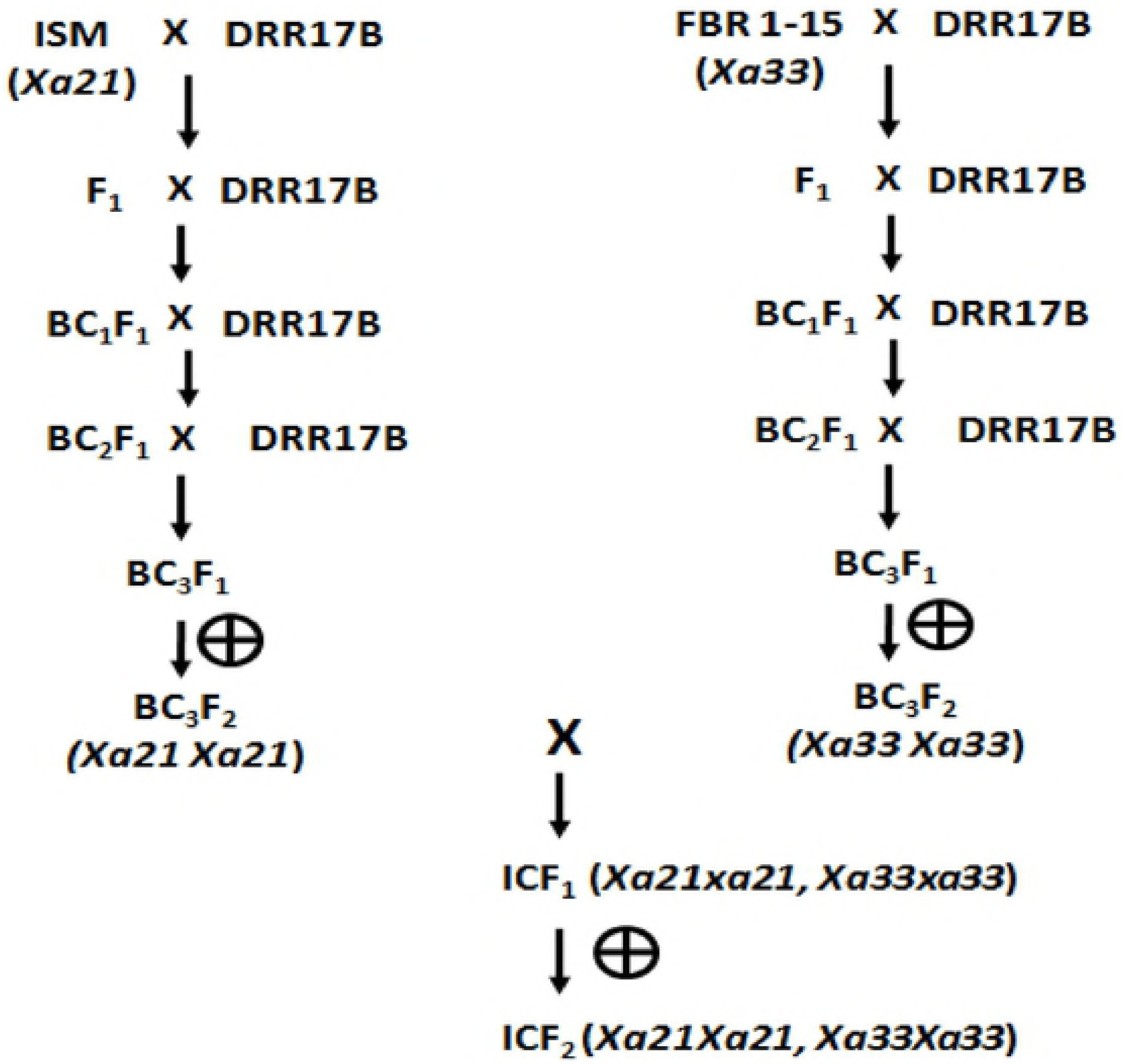
Marker-assisted backcrossing scheme adopted in the study.

The process of marker-assisted backcrossing was repeated until BC_3_ generation, and BC_3_F_1_ plants of DRR17B possessing either *Xa21* or *Xa33* and maximum recovery of recurrent parent genome were then selfed to obtain BC_3_F_2_s. Plants homozygous for either *Xa21* or *Xa33* were identified among the BC_3_F_2_ plants and the best plants from the two crosses were intercrossed to obtain intercross F_1_s (i.e., ICF_1_s). ‘True’ ICF_1_ plants were identified by screening with molecular markers specific for *Xa21* and *Xa33* and were then selfed to generate intercross F_2_s (i.e., ICF_2_s). Plants homozygous for both *Xa21* and *Xa33* were identified among the ICF_2_ plants using the gene-specific markers. The identified plants were advanced through the pedigree method of selection (involving selfing followed by morphological trait-based visual selection) up to ICF_4_ generation. Marker-assisted selection procedures were followed as recommended by [16] and [17] for detection of *Xa21* and *Xa33* genes, while the protocol described by [18] was adopted for background selection and detection of non-restoring alleles of *Rf4* and *Rf3*.

### Screening for BB resistance

Eight virulent isolates of the BB pathogen, *Xanthomonas oryzae* pv. *oryzae* (*Xoo*) collected from BB disease endemic locations in India, *viz*., IX-020 (Hyderabad, Telangana), IX-133 (Raipur, Chhattisgarh), IX-027 (Chinsurah, West Bengal), IX-200 (Pantnagar, Uttarakhand), IX-002 (Faizabad, Uttar Pradesh), IX-403 (Thanjavur, Tamil Nadu), IX-090 (Ludhiana, Punjab) and IX-281 (Tanuku, Andhra Pradesh) were used to screen the ICF_4_ lines of DRR17B (possessing the gene combinations *Xa21*+*Xa33, Xa21* alone or *Xa33* alone) along with the donor parents/resistant check, ISM (possessing *xa5*+*xa13*+*Xa21*), FBR1-15 (possessing *Xa33*) and BB recurrent parent and susceptible check (DRR17B and TN1) were screened under glasshouse conditions for their resistance/susceptibility against BB. The *Xoo* strains were cultured and stored as described by [12]. The rice plants were clip-inoculated with a bacterial suspension of 10^8-9^ CFU/ml at maximum tillering stage (45 to 55 days after transplanting) through the methodology of [20]. Approximately, 5 to 10 leaves were inoculated per plant, and the disease reaction was scored 14 days after inoculation [21]. In addition to measurement of BB lesion length, the disease score was calculated as per IRRI Standard Evaluation System (SES) scale, which is based on percent diseased leaf area [22].

### Screening for agro-morphological characters

Improved lines of DRR17B (ICF_4_) were field evaluated in randomized complete block design in kharif 2014 for the agro-morphological traits involving days to 50% flowering (days), plant height (cm), number of productive tillers (No.), panicle length (cm), grains per panicle (No.) and spikelet fertility. Each entry was planted in 20 rows of 4m length with a spacing of 15 x 20 cm between rows and within rows. Days to 50 percent flowering was recorded based on number of days from sowing to 50% population flowering on a whole plot basis. Plant height (cm), number of productive tillers (No.) and panicle length (cm) were recorded from 5 competitive plants from each plot chosen at random and the mean values computed for different lines. Five individual panicles harvested separately from five plants were collected to compute for the average grain number per panicle (No.). The Improved lines were crossed with IR58025A line and evaluated for spikelet fertility based on seed setting of each cross. The percentage was calculated based on seed setting per panicle as described in [18].

### Statistical analysis

Agro-morphological and phenotypic BB screening data were analysed using standard procedures by [23]. Analysis of variance (ANOVA) and Least Significance Difference (LSD) at 5% significant standard error of Mean (S.E.M ±) were calculated by using MS Excel and Statistix 8.1 software to explore the variation between improved lines and parents.

## Results

### Marker-Assisted Transfer of *Xa21* and *Xa33* into DRR17B

The true F_1_s derived by crossing DRR17B with ISM (i.e., Cross I) and FBR1-15 (i.e., Cross II) were backcrossed with DRR17B to obtain BC_1_F_1_s, which were then screened with the gene-specific markers. A total of 61 and 65 BC_1_F_1_ plants were observed to be positive for the target genes in Cross I and Cross II, respectively. The positive plants were screened with markers specific for *Rf3* and *Rf4*, and a total of 15 and 11 plants were identified to be devoid of both the fertility restorer genes concerning Cross I and Cross II, respectively. These plants were then subjected to background selection using a set of polymorphic SSR markers (61 markers for Cross I and 64 for Cross II). Plant # IIRRGP3 from Cross I, with a recurrent parent genome (RPG) recovery of 73.7% and Plant # IIRRGP22 from Cross II, with a RPG recovery of 75% were identified to be the best ones (i.e. having a maximum recovery of DRR17B genome) and were used for further backcrossing. The process of marker-assisted backcrossing was carried out until BC_3_F_1_ generation (details given in Table 1). At BC_3_F_1_, plant # IIRRGP3-87-64 from Cross I with RPG recovery of 93.4% and plant # IIRGP22-73-10 with RPG recovery of 93.7% were identified to be superior and were selfed to generate BC_3_F_2_s. With regards to the BC_3_F_2_s produced from Cross I and Cross II, 39 and 52 plants were identified to be homozygous for *Xa21* and *Xa33*, respectively. Among these, a solitary plant, which was morphologically similar to DRR17B, was identified from Cross I (i.e., plant # IIRRGP 3-87-64-22 and Cross II (i.e., plant # IIRRGP 22-73-10-15) and intercrossed with each other to generate intercross F_1_s (i.e., ICF_1_s). Out of 68 ICF_1_s, 63 were identified to be heterozygous for both *Xa21* and *Xa33* (i.e. true intercross F_1_s), and they were selfed to obtain ICF_2_ generation. At ICF_2_, a total of 309 plants were screened with markers specific for *Xa21* and *Xa33* and 18 were identified to be double homozygotes (Table 1; Fig 2). A total of nine plants out of the 18, which were identified to be phenotypically similar to DRR17B, were further advanced until ICF_4_ generation through phenotype-based pedigree selection. At ICF_4_ generation, six promising lines which were similar to the recurrent parent were identified (line #IIRRIC 10-8-94, IIRRIC 10-19-138, IIRRIC 102-26-7, IIRRIC 123-34-84, IIRRIC 123-58-3 and IIRRIC 172-77-12) and analysed for their resistance to BB, sterility maintenance ability and also characterized for important agro-morphological traits. Among the six improved lines, line # IIRRIC102-26-7 exhibited the highest recurrent parent genome recovery with more than 95% along with minimal linkage drag on carrier chromosomes (Fig 3).

**Fig 2.**
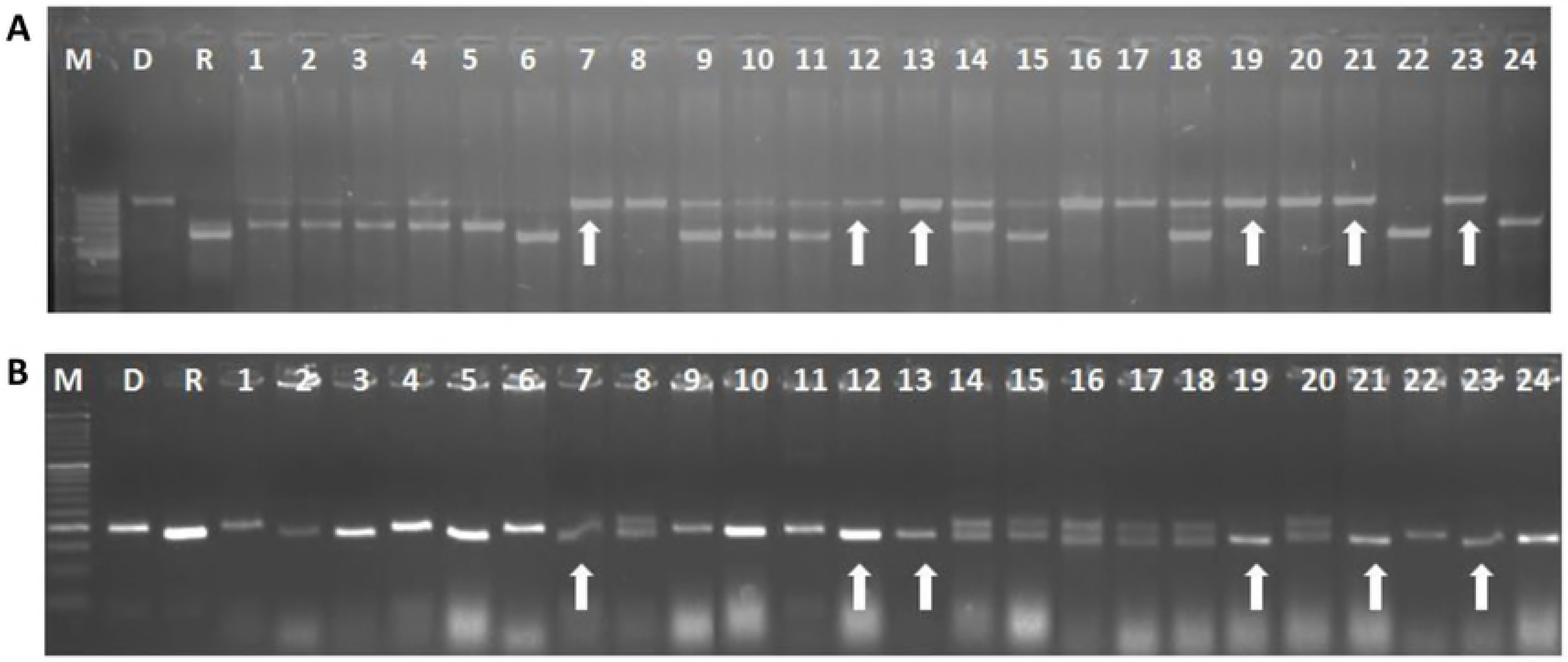
Screening of ICF_2_ population for identification of double homozygotes for the target resistance genes, viz., *Xa21* and *Xa33*. The ICF_2_ plants were screened through PCR to analyze the allelic status of *Xa21* (A) and *Xa33*(B) using the gene-specific markers. M – Marker, R – Recurrent parent (i.e. DRR17B) and D– donor parent [i.e., ISM (A) and FBR1-15EM (B)]. Arrows indicate plants which possess target genes *Xa21* and *Xa33* in homozygous condition.

**Fig 3.**
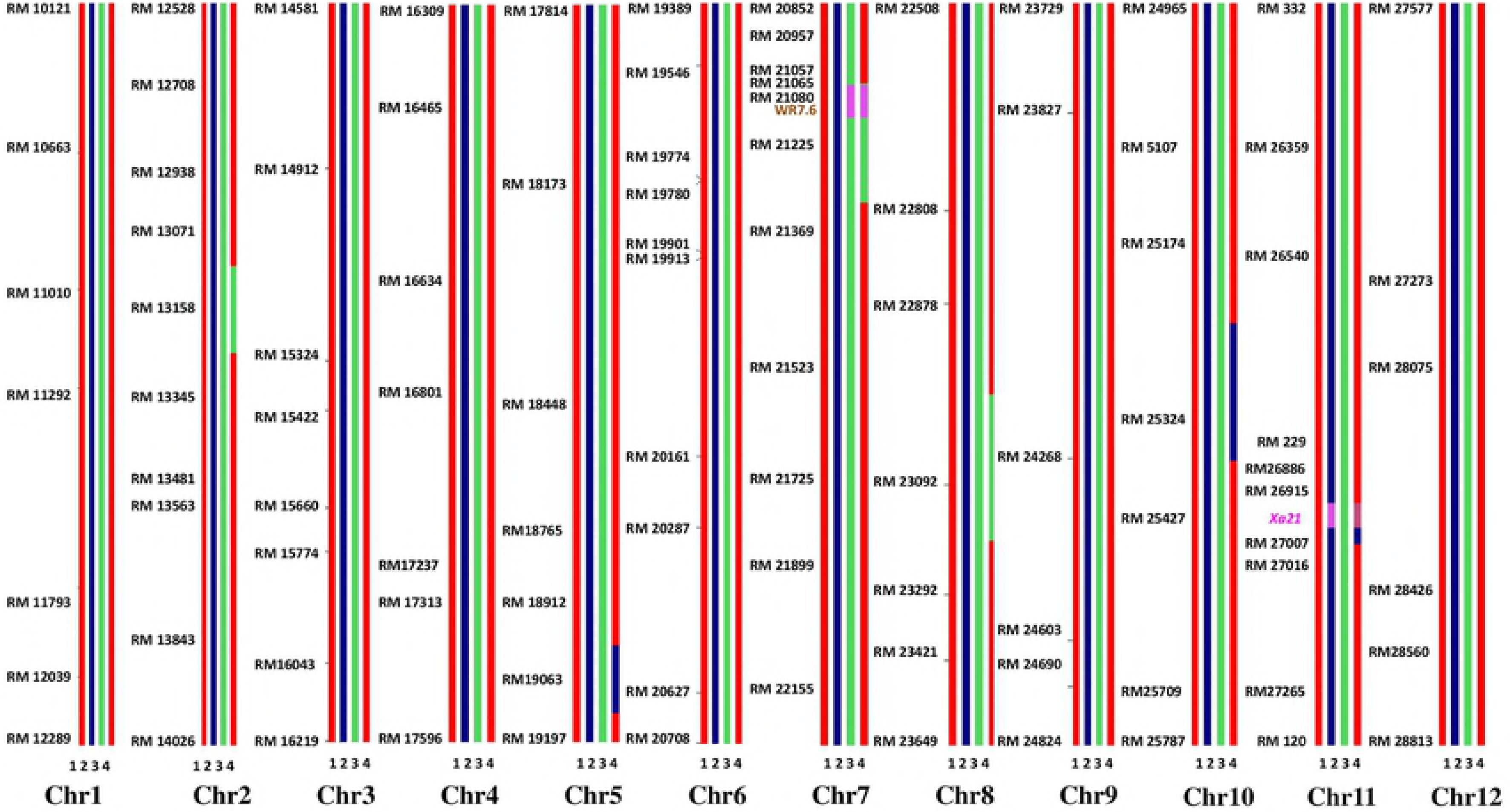
Graphical genotyping representation for the best improved line (IIRRIC102-26-7) of DRR17B. Graphical genotyping representing that the best line of improved two-gene-containing DRR17B line (*Xa21*+ *Xa33*), IIRRIC102-26-7 exhibiting the highest genome recovery of the recurrent parent with more than 95%, along with minimum linkage drag on carrier chromosomes 7 and 11, with less than 2 Mb donor parent chromosome (1 DRR17B, 2 ISM, 3 FBR1-15, 4 IIRRIC102-26-7).

**Table 1.**
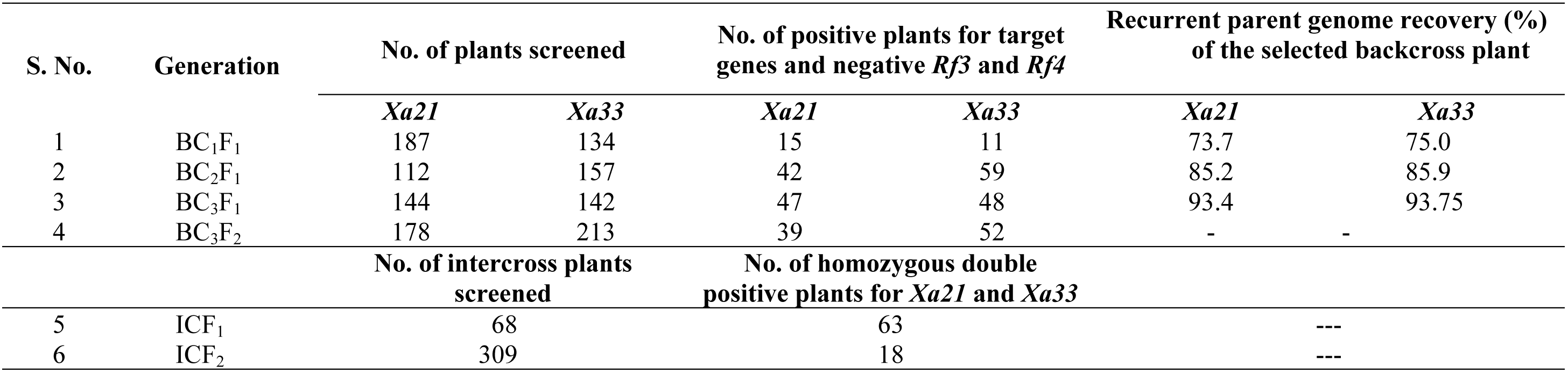
Details of plants generated and analyzed with markers in each generation of backcrossing/intercrossing.

### Phenotypic evaluation of improved lines for BB resistance

The recurrent parent, DRR17B (with lesion lengths ranging from 18.8 to 33.1 cm) and susceptible check TN1 (with lesion lengths ranging from 20.9 to 33.8 cm) showed a disease score of 9 against all the eight isolates of the *Xoo* (Table 2; Fig 4 depicted as graph). The resistant check and the donor for *Xa21* gene ISM (possessing *Xa21, xa13*, and *xa5*) showed a score of 3 against all the isolates (with an average lesion length ranging from 1.6 to 3.6 cm). FBR1-15, the donor for *Xa33* gene and improved DRR17B lines possessing *Xa33* (# IIRRGP22-73-10-15-13-2) showed a resistance score of 3 with most of the isolates (with lesion lengths ranging from 1.7 to 4.8 cm), with two isolates, IX-002 and IX-281 recorded moderate resistance reaction with a score of 5 (average lesion lengths ranging from 7.3 to 9.7 cm and 7.5 to 9.2 cm). The improved lines of DRR17B containing only *Xa21* (# IIRRGP3-87-64- 22-4-50) showed a resistance reaction against two isolates *viz*., IX-002 and IX-090 with a score of 3 (with lesion lengths of 2.8 to 4.3 cm and 2.0 to 2.8 cm, respectively), while with three isolates, *viz*., IX-020, IX-027 and IX-281, the line with only *Xa21* exhibited moderately susceptibility with a score of 7 (with lesion lengths of 12.5 to 14.7 cm, 13.1 to 14.5 cm and 13.0 to 14.6 cm, respectively). Further, the line showed highly susceptible reaction with a score of 9 (with lesion lengths of 20.1 to 23.5cm, 22.2 to 25.6 cm and 21.9 to 24.4cm, respectively) with three otherr three isolates *viz*., IX-133, IX-200 and IX-409, respectively. The improved lines of DRR17B containing both *Xa21* + *Xa33* exhibited a significantly higher level of resistance, showing a score of 1 against all eight isolates with lesion lengths ranging from 0.1 to 1 cm (Table 2; Fig 4).

**Fig 4.**
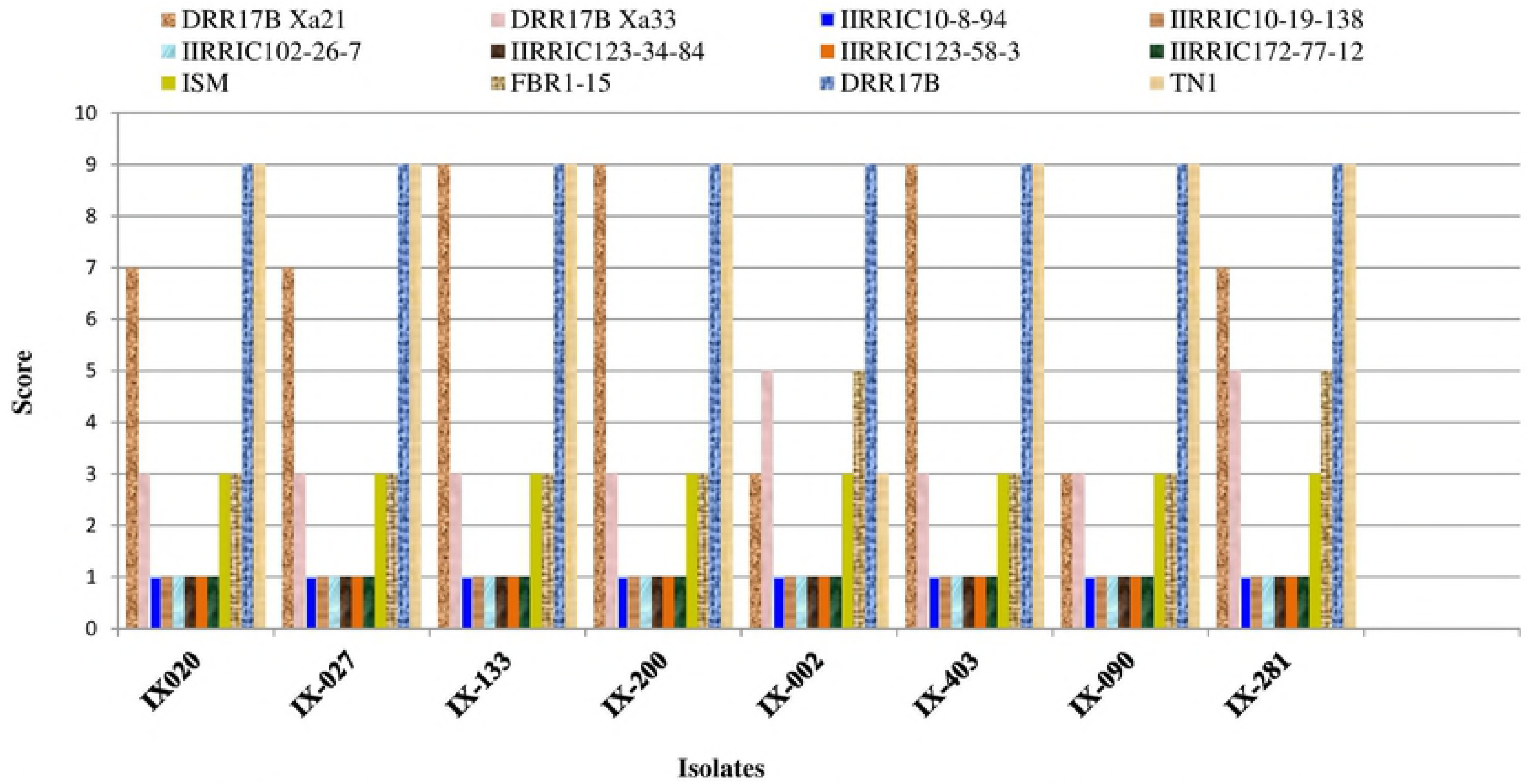
Screening of the single-gene and two-gene pyramid lines of DRR17B against different virulent isolates of the bacterial blight pathogen. Eight selected ICF_4_ of DRR17B possessing either *Xa21 (*IIRRGP3-87-64-22-4-50) or *Xa33* (IIRRGP22-73-10-15-13-2) or Xa21 + *Xa33* (# IIRRIC10-8-94, IIRRIC10-19-138, IIRRIC102-26-7, IIRRIC123-34-84, IIRRIC123-58-3, and IIRRIC172-77-12) were screened for their BB resistance with eight virulent isolates of *Xanthomonas oryzae* pv. *oryzae* (*Xoo*) along with the recurrent parent (DRR17B) and donor parents (ISM and FBR1-15). While all the lines showed excellent resistance against the multiple isolates of *Xoo* screened, the two-gene pyramid lines (i.e., *Xa21 + Xa33*) were observed to show a higher level of resistance to the different isolates of the pathogen.

**Table 2.** Reaction of the breeding lines of DRR17B possessing *Xa21* and *Xa33*, singly or in combination against eight virulent isolates of the bacterial blight pathogen, *Xanthomonas oryzae* pv. *oryzae (Xoo)*.

| *Xoo* Isolates |  | Breeding Lines |  |  |  |  |  |  |  | Parents/Resistant/Susceptible Check |  |  |  | CV | LS<br>D | H<sup>2</sup> | F<br>Value |
| --- | --- | --- | --- | --- | --- | --- | --- | --- | --- | --- | --- | --- | --- | --- | --- | --- | --- |
|  |  | 1 | 2 | 3 | 4 | 5 | 6 | 7 | 8 | 9 | 10 | 11 | 12 |  |  |  |  |
| IX-020 | Mean ± | 13.66 ± | 3.8 ± | 0.68 ± | 0.5 ± | 0.58 ± | 0.66 ± | 0.54 ± | 0.56 ± | 2.66 ± | 3.68 ± | 31.02 ± | 30.12 ± | 11.5 | 1.08 | 0.99 | 909.86 |
|  | SE | 0.42 b | 0.28c | 0.07e | 0.06e | 0.06e | 0.07e | 0.05e | 0.05e | 0.19d | 0.27cd | 0.87a | 0.95a |  |  |  |  |
|  | Range | 12.5-14.7 | 3.2-4.7 | 0.5-0.9 | 0.3-0.7 | 0.4-0.7 | 0.5-0.7 | 0.4-0.7 | 0.5-0.7 | 2.1-3.1 | 3.0-4.4 | 28.6-33.1 | 27.6-32.8 |  |  |  |  |
|  | Score | 7 | 3 | 1 | 1 | 1 | 1 | 1 | 1 | 3 | 3 | 9 | 9 |  |  |  |  |
| IX-027 | Mean ± | 13.88 ± | 4.08 ± | 0.28 ± | 0.44 ± | 0.42 ± | 0.34 ± | 0.30± | 0.36 ± | 2.90 ± | 3.94 ± | 21.60 ± | 25.20 ± | 17.8 | 1.39 | 0.99 | 335.00 |
|  | SE | 0.24c | 0.24d | 0.10e | 0.07e | 0.07e | 0.06e | 0.09e | 0.09e | 0.23d | 0.30d | 0.90b | 1.30a |  |  |  |  |
|  | Range | 13.1-14.5 | 3.5-4.9 | 0.1-0.6 | 0.2-0.6 | 0.2-0.6 | 0.2-0.5 | 0.1-0.6 | 0.1-0.6 | 2.1-3.5 | 3.1-4.8 | 18.8-23.5 | 20.9-28.5 |  |  |  |  |
|  | Score | 7 | 3 | 1 | 1 | 1 | 1 | 1 | 1 | 3 | 3 | 9 | 9 |  |  |  |  |
| IX-133 | Mean ± | 22.04 ± | 3.56 ± | 0.2 ± | 0.24 ± | 0.26 ± | 0.26± | 0.28 ± | 0.22 ± | 2.2 ± | 3.4 ± | 24.14 ± | 26.06 ± | 11.91 | 1.04 | 0.99 | 809.23 |
|  | SE | 0.79c | 0.25d | 0.00f | 0.02f | 0.04f | 0.04f | 0.04f | 0.02f | 0.19e | 0.21d | 0.75b | 0.57a |  |  |  |  |
|  | Range | 20.1-23.5 | 2.7-4.1 | 0.2 | 0.2-0.3 | 0.2-0.4 | 0.2-0.4 | 0.2-0.4 | 0.2-0.3 | 1.7-2.7 | 2.8-3.9 | 22.9-26.5 | 24.8-27.5 |  |  |  |  |
|  | Score | 9 | 3 | 1 | 1 | 1 | 1 | 1 | 1 | 3 | 3 | 9 | 9 |  |  |  |  |
| IX-200 | Mean ± | 23.42 | 3.52 ± | 0.82 ± | 0.78 ± | 0.7± | 0.76 | 0.68 ± | 0.64 | 2.74 ± | 3.68 ± | 30.56 ± | 31.62 ± | 10.49 | 1.11 | 0.99 | 1006.00 |
|  | SE | ±0.70b | 0.21c | 0.08d | 0.04d | 0.07d | ±0.08d | 0.08d | ±0.07d | 0.18c | 0.17c | 0.76a | 0.79a |  |  |  |  |
|  | Range | 22.2-25.6 | 3.0-4.2 | 0.6-1 | 0.7-0.9 | 0.5-0.9 | 0.5-1 | 0.4-0.8 | 0.4-0.8 | 2.3-3.1 | 3.3-4.2 | 28.2-32.8 | 29.3-33.8 |  |  |  |  |
|  | Score | 9 | 3 | 1 | 1 | 1 | 1 | 1 | 1 | 3 | 3 | 9 | 9 |  |  |  |  |
| IX-002 | Mean ± | 3.72 ± | 8.58 ± | 0.86 ± | 0.74 ± | 0.8 | 0.72 ± | 0.84 ± | 0.8 ± | 2.34 ± | 8.72 ± | 24.88 ± | 24.76 ± | 13.22 | 1.09 | 0.99 | 558.46 |
|  | SE | 0.26c | 0.49b | 0.05e | 0.07e | ±0.04e | 0.06e | 0.05e | 0.04e | 0.22d | 0.43b | 0.90a | 0.65a |  |  |  |  |
|  | Range | 2.8 - 4.3 | 7.2-9.8 | 0.7-1 | 0.5-0.9 | 0.7-0.9 | 0.6-0.9 | 0.7-1 | 0.7-0.9 | 1.8-2.7 | 7.3-9.7 | 22.6-27.7 | 22.5-26.2 |  |  |  |  |
|  | Score | 3 | 5 | 1 | 1 | 1 | 1 | 1 | 1 | 3 | 5 | 9 | 3 |  |  |  |  |
| IX-403 | Mean ± | 23.34 ± | 2.52 | 0.48 ± | 0.52 | 0.48 ± | 0.5 ± | 0.48 | 0.52 ± | 2.28 ± | 2.84 ± | 21.34 ± | 25.92± | 13.21 | 1.19 | 0.99 | 658.00 |
|  | SE | 0.53b | ±0.15c | 0.06d | ±0.07d | 0.06d | 0.07d | ±0.06d | 0.02d | 0.19c | 0.17c | 0.83a | 0.98a |  |  |  |  |
|  | Range | 21.9-24.4 | 2-2.7 | 0.3-0.6 | 0.3-0.7 | 0.4-0.7 | 0.3-0.7 | 0.3-0.6 | 0.5-0.6 | 1.6-2.7 | 2.4-3.3 | 23.9-28.1 | 23.4-28.8 |  |  |  |  |
|  | Score | 9 | 3 | 1 | 1 | 1 | 1 | 1 | 1 | 3 | 3 | 9 | 9 |  |  |  |  |
| IX-090 | Mean ± | 2.46 ± | 2.88 | 0.56 ± | 0.56 ± | 0.54 ± | 0.58 ± | 0.54 | 0.6 ± | 1.96 ± | 3.22 ± | 25.78 ± | 29.42 ± | 12.03 | 0.88 | 0.99 | 1101.20 |
|  | SE | 0.13cd | ±0.17c | 0.04e | 0.06e | 0.07e | 0.06e | ±0.04e | 0.04e | 0.26d | 0.19c | 0.74b | 0.66a |  |  |  |  |
|  | Range | 2-2.8 | 2.4-3.5 | 0.5-0.7 | 0.4-0.7 | 0.3-0.7 | 0.4-0.7 | 0.5-0.7 | 0.5-0.7 | 1.7-2.8 | 1.7-3.7 | 20.9-28.5 | 23.2-31.6 |  |  |  |  |
|  | Score | 3 | 3 | 1 | 1 | 1 | 1 | 1 | 1 | 3 | 3 | 9 | 9 |  |  |  |  |
| IX-281 | Mean ± | 13.68 ± | 8.3 ± | 0.54 ± | 0.56 ± | 0.62 ± | 0.66 ± | 0.46 ± | 0.6 ± | 2.92 ± | 8.24 | 24.1 ± | 28.68 ± | 9.43 | 1.53 | 0.99 | 987.94 |
|  | SE | 0.40c | 0.45d | 0.09f | 0.04f | 0.08f | 0.05f | 0.05f | 0.04f | 0.22e | ±0.30d | 0.56b | 0.55a |  |  |  |  |
|  | Range | 13.0 - 14.6 | 7.1 - 9.4 | 0.3- 0.8 | 0.5-0.7 | 0.4-0.8 | 0.5-0.8 | 0.3-0.6 | 0.5-0.7 | 2.4-3.6 | 7.5-9.2 | 23.5-25.7 | 27.9-30.3 |  |  |  |  |
|  | Score | 7 | 5 | 1 | 1 | 1 | 1 | 1 | 1 | 3 | 5 | 9 | 9 |  |  |  |  |

Breeding line **1** represents DRR17B line containing *Xa21* gene that was screened with eight different isolates under glass house conditions at IIRR. With two isolates (*viz*., FZB and Lud-05-1 *Xa21*) introgressed lines that showed resistance reaction having a score of 3. With remaining isolates, *Xa21* introgressed lines showed moderate susceptibility with a score of 7 and high susceptibility having a score of 9 (Lore et al., 2011). Breeding line **2** represents DRR17B line containing *Xa33* gene, except for two isolates (*viz*., FZB and TNK12-3 moderate resistance with a score of 5) while the remaining isolates showed a resistance reaction score of 3. Breeding lines **3-8** represent improved lines of DRR17B containing *Xa21* + *Xa33* genes that exhibited a high level of resistance with a score of 1 against all eight isolates. Resistant checks **9** and **10** represent ISM, which showed resistance against all eight isolates with a score of 3 and another resistant check, FBR1-15, which showed resistance reaction score of 3 except the two isolates that showed moderate resistance with a score of 5. Recurrent parents **11** and **12** represent DRR17B and susceptible check TN1 that exhibited highly susceptible reactions against all eight isolates with a score of 9. (**1**-IIRRGP3-87-64-22-4-50 (*Xa21*), **2**-IIRRGP22-73-10-15-13-2 (*Xa33*), **3**-IIRRIC10-8-94, **4**-IIRRIC10-19-138, **5**-IIRRIC102-26-7, **6**-IIRRIC123-34-84, **7**-IIRRIC123-58-3, **8**-IIRRIC172-77-12, **9**-ISM, **10**-FBR1-15, **11**-DRR17B and **12**-TN1).

### Characterization of ILs for maintainance ability and agro-morphological traits

We screened the six improved lines for their maintainance ability. Three showed partial spikelet fertility, while the remaining three lines (viz., line # IIRRIC102-26-7, IIRRIC123-34-84, and IIRRIC172-77-12) showed complete spikelet sterility when crossed with the WA-CMS line, IR58025A (Table 3).With regards to five agro-morphological parameters (days to 50% flowering, plant height, number of productive tillers, panicle length and number of grains per panicle), all the six improved lines were identical in their panicle length and number of productive tillers to DRR17B, while differences were observed with respect to the number of grains per panicle. The improved lines viz., IIRRIC10-8-94, IIRRIC102-26-7, IIRRIC123-58-3 and IIRRIC172-77-12 possessed more number of grains per panicle-301.6, 360.4, 308 and 317, respectively (Figs 5A and 5B). However, concerning plant height, all the six lines showed shorter plant height as compared to DRR17B. As with regards to panicle length, line # IIRRIC102-26-7 was observed to have longer panicles with an average length of 24.16 cm, the remaining improved lines exhibited equal or less than the recurrent parent DRR17B (average length of 23.24 cm: Table 3). Line # IIRRIC102-26-7 displayed more numbers of productive tillers per plant (average of 12), all remaining improved lines were similar to the recurrent parent. With regards to the days to 50% flowering, all the six improved lines flowered earlier (between 92-102 days), as compared to DRR17B, which took 105 days.

**Fig 5A.**
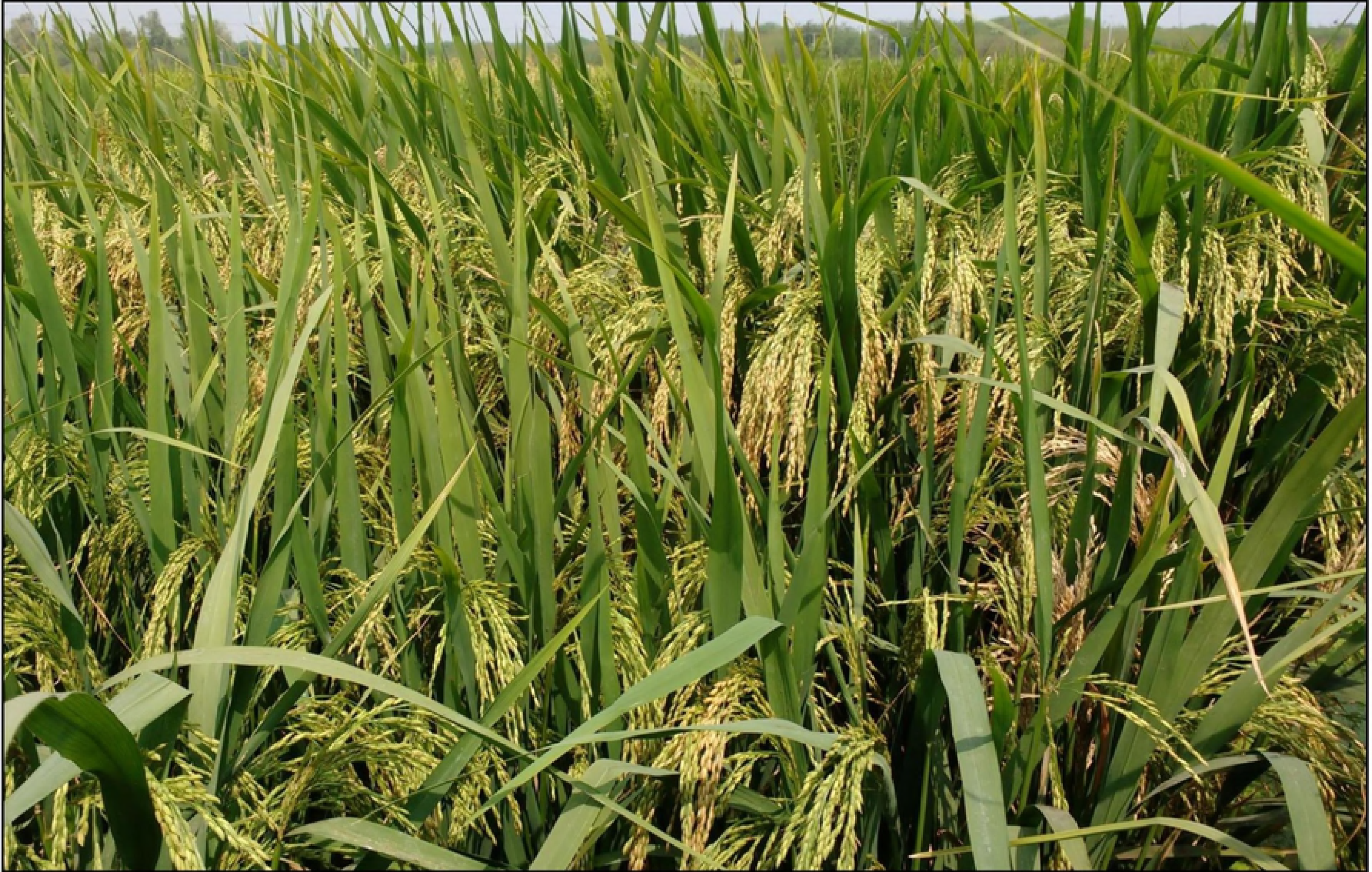
Improved line of DRR17B (# IRIC102-26-27) displaying high grain number under field conditions at IIRR.

**Fig 5B.**
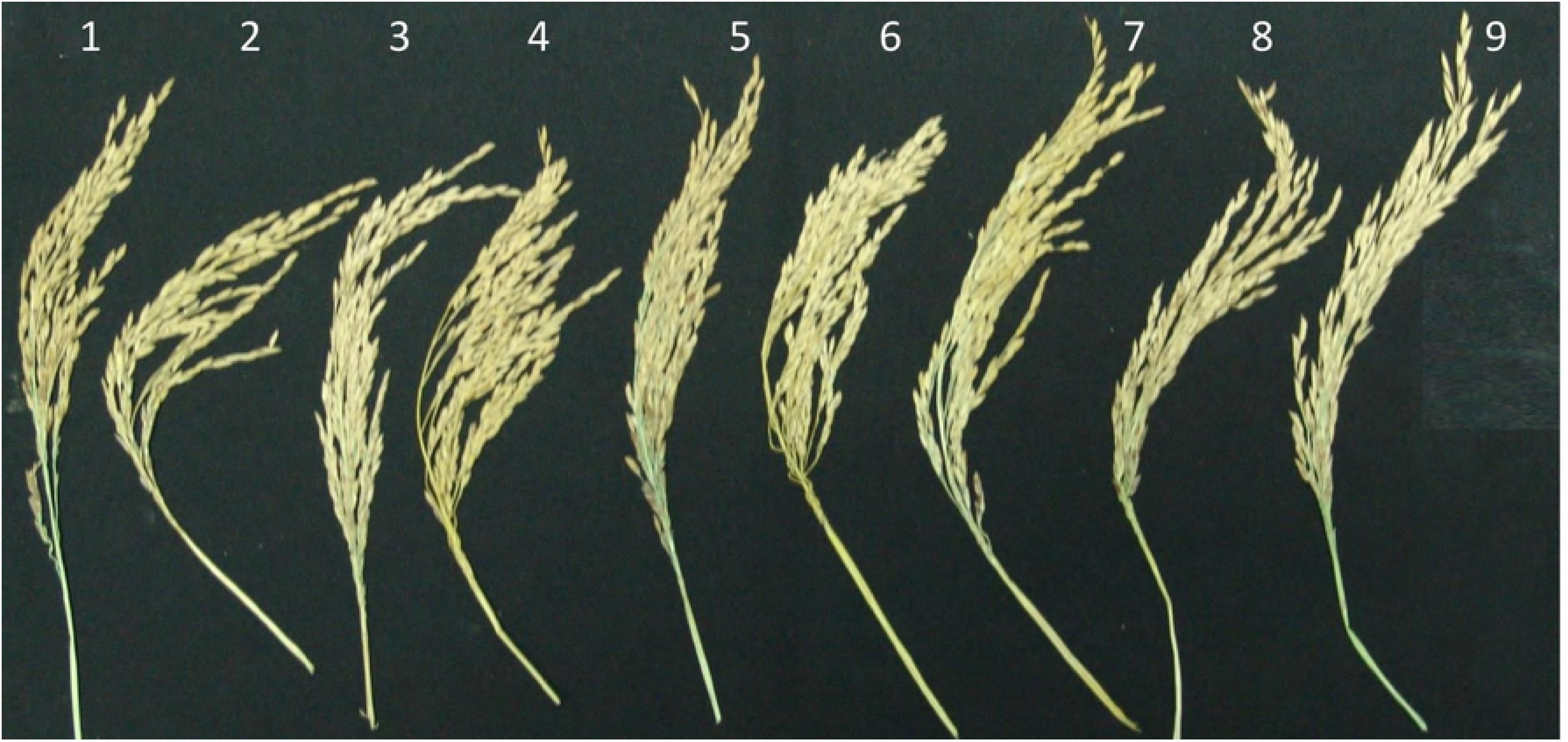
Panicles of improved DRR17B lines along with donor and recurrent parents. The improved lines of DRR17B possessing *Xa21* + Xa33 genes (# IIRRIC10-8-94, IIRRIC10-19-138, IIRRIC102-26-7, IIRRIC123-34-84, IIRRIC123-58-3, and IIRRIC172-77-12) were displayed more number of grains per panicles and similar panicle length (except number 6) when compared with DRR17B

**Table 3.**
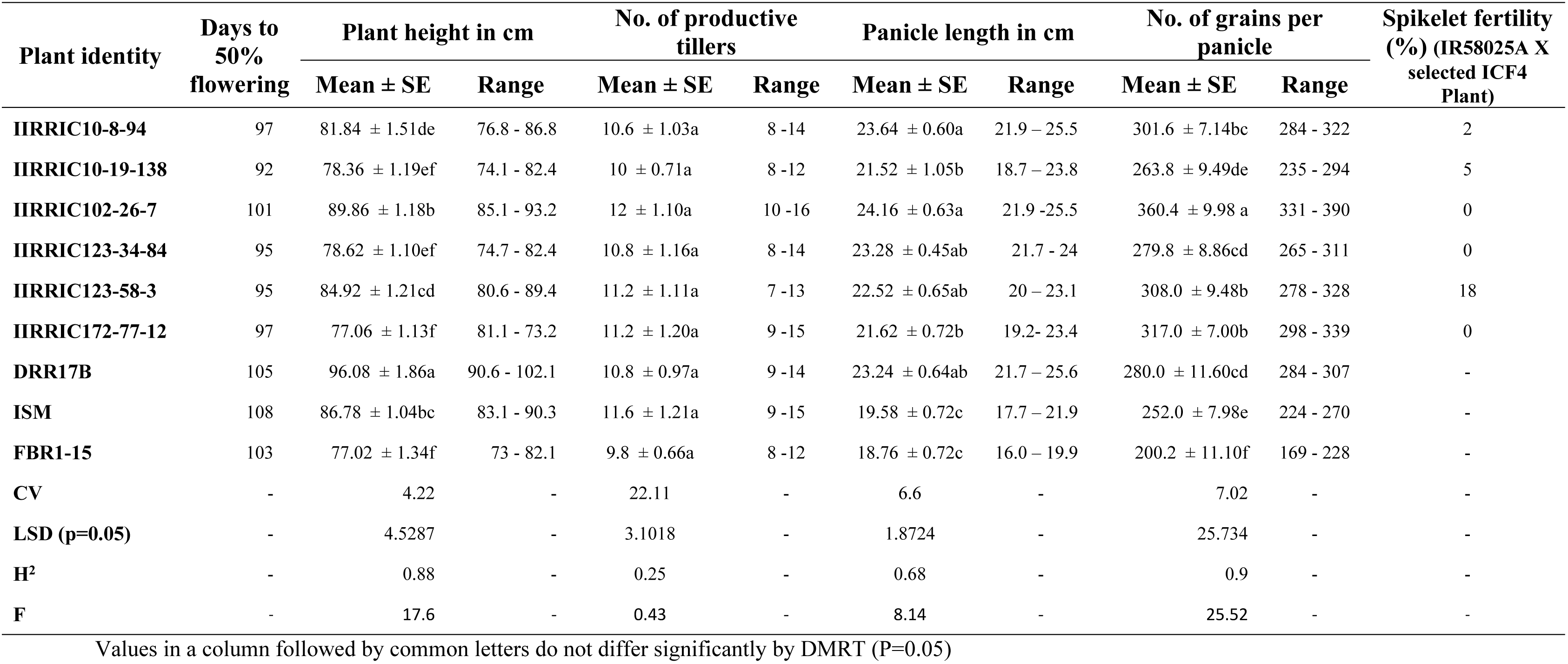
Agro-morphological features of selected backcross derived lines of DRR17B possessing *Xa21 + Xa33*.

## Discussion

Several studies indicate that global rice production needs to be doubled by 2050 to meet the demands of ever growing population [24]. However, rice grain yield is badly affected by biotic and abiotic stresses including diseases and pests [25]. The present study was taken up to improve, an elite maintainer of rice, DRR17B, for its resistance against BB resistance. DRR17B is a fine grain type, medium duration, maintainer line of rice, possessing stable maintenance ability and was developed by ICAR-Indian Institute of Rice Research, Hyderabad, India [18]. As DRR17A and its maintainer parent-DRR17B are highly susceptible to BB disease, this study was carried out with an objective to introgress two major dominant BB resistant genes, *viz*., *Xa21* and *Xa33* through MABB in order the make the maintainer line durably resistant to BB. The two selected genes are known to confer resistance against multiple isolates of the BB pathogens; hence, the hybrids developed from improved lines of DRR17A will also be durably resistant to the disease.

Introgression of disease resistance genes through conventional breeding involving morphology-based phenotypic selection is a laborious, time consuming process and its success significantly depends on the existence of environmental conditions favourable for disease development and the availability of appropriate virulent strains of the pathogen causing the disease [9]. As compared to conventional breeding, marker-assisted selection (MAS) is a simpler strategy for targeted introgression of resistance genes as it does not depend on the availability of virulent strains or existence of ideal environmental conditions, since the selections are indirect, and are based on the presence or absence of specific alleles of molecular markers linked to the resistance genes. Earlier [18], [26] and [27] successfully developed bacterial blight resistant versions of hybrid rice parental lines PRR78 and IR58025B, through marker-assisted selection for target traits in the initial stages and phenotype-based selection at later stages and the same methodology was also adopted in the current study.

So far, at least 41 genes conferring resistance against BB have been identified in rice [9, 10, 11]. Among them, the wild rice derived gene, *Xa21* encoding a receptor kinase-like protein has been successfully deployed by many research groups across the world, as it has been documented to confer broad-spectrum resistance against the BB disease [13, 14, 18, 26, 28, 29, 30, 31, 32]. The commonly used BB resistance gene *Xa21* has been tagged and mapped on chromosome 11 with a tightly-linked PCR-based marker pTA248 [16]. *Xa33*, the wild rice derived BB resistance gene has been reported to confer broad spectrum resistance [17] and the gene has been deployed by the research group at Tamil Nadu Agricultural University, Coimbatore, India and the breeding lines possessing *Xa33* were observed to be very effective in terms of their BB resistance [33, 34]. Hence, these two broad spectrum resistance genes were selected for introgression into the DRR17B.

Phenotypic screening for BB resistance was carried out in this study among selected single gene containing BC_3_F_6_ lines possessing either *Xa21* or *Xa33* and two-gene containing intercross derived lines at ICF_4_ generation possessing *Xa21*+*Xa33* along with the donor and recurrent parents (ISM, FBR1-15, and DRR17B, respectively) using eight virulent isolates of *Xoo*. All the improved lines possessing *Xa21+Xa33* were observed to show significantly higher level of resistance against BB when compared to the donor parents, ISM and FBR1-15. Single gene containing lines of DRR17B (i.e. possessing either *Xa21* or *Xa33*), the recurrent parent DRR17B and the BB susceptible check TN1 (Table 2; Fig 4). It is earlier known that *Xa21* confers broad spectrum resistance against many of the virulent pathotypes of *Xoo* in India [13, 14] and several studies have indicated the suitability of *Xa21* in BB resistance gene pyramiding programmes [8, 14, 28, 35, 36]. However, in this study, a few isolates of the pathogen were observed to be compatible with *Xa21* containing lines of DRR17B indicating that *Xoo* isolates, which are capable of overcoming *Xa21* conferred resistance are fast-developing [13, 37, 38]. Interestingly, the improved lines of DRR17B possessing *Xa33* were observed to show a better level of resistance as compared to the lines having *Xa21.* Furthermore, DRR17B lines possessing both *Xa21* and *Xa33* were observed to be highly resistant against all the eight virulent isolates of *Xoo*, thus, indicating the suitability of deployment of *Xa33* either singly or in combination with *Xa21.* Earlier, two elite restorer lines, KMR3R, and RPHR1005 were improved for BB resistance by introducing *Xa21* [18, 27, 30, 32]. Similarly, *Xa33* has been deployed in different genetic backgrounds by different research groups [15, 17, 33, 34]. However, this is the first report wherein *Xa21* has been combined with *Xa33* in the genetic background of an elite maintainer line, i.e., DRR17B and the gene-pyramid lines demonstrated a higher level of resistance as compared to lines possessing a single resistance gene (Table 2; Fig 4).

It is a known fact that long term cultivation of rice varieties possessing single resistance gene can result in the breakdown of resistance by faster development of virulent pathogens [37, 38, 39]. Hence, pyramiding of multiple resistance genes has been advocated to be one of the best strategies to achieve durable dual-resistance [14, 40, 41]. In our present study, the genotype ISM (with *Xa21* + *xa13* + *xa5*) has displayed satisfactory level of resistance with a score of 3 against all eight isolates. Interestingly, when *Xa21* gene was combined with another major dominant gene- *Xa33*, such breeding lines exhibited the highest level of resistance with a score of 1. This indicates that the gene combination *Xa21 + Xa33* displayed a broad spectrum of resistance and hence can be recommended for deployment in hybrid rice improvement programs as both *Xa21* and *Xa33* are both dominant and will express in the F_1_ hybrid.

Similar to the approach adopted in the current study, several earlier studies also resorted to phenotype-based selection for key agro-morphological traits along with marker-assisted selection while improving elite varieties and parental lines for one or more target traits through MABB [14, 18, 27, 29, 30, 31, 32, 42]. The approach of deployment of MABB strategy for the target resistance genes along with negative selection for major fertility restorer genes, *Rf3* and *Rf4*, coupled with phenotype-based selection for certain key agronomic characters helped in near-complete recovery of good features of DRR17B along with identification of few improved lines with complete maintenance ability (Table 3). In addition to improving BB resistance of DRR17B, we also focused on the identification of improved lines of DRR17B possessing plant height which is significantly shorter than DRR17B, as shorter plant is preferred for deployment as good maintainers [18]. Significant differences in plant height were observed in many improved DRR17B lines *viz*., RMSIC 10-8-94, RMSIC 10-19-138, RMSIC 102-26-7, RMSIC 123-34-84, RMSIC 123-58-3 and RMSIC 172-77-12 and these lines could serve as better maintainers as compared to DRR17B. Interestingly, when compared to DRR17B, some of the improved lines exhibited advantage concerning grain number per panicle. These lines include RMSIC 10-8-94, RMSIC 102-26-7, RMSIC 123-58-3 and RMSIC 172-77-12 (Figs 5A and 5B). For the panicle length, all the improved lines showed values equivalent to DRR17B, except one line *viz*., RMSIC 102-26-7, a wherein slight improvement over the recurrent parent was noticed. Significant differences (i.e., shorter duration) were observed concerning number of days to 50% flowering in some of the backcross derived plants (Table 3). No significant differences were observed between improved versions of DRR17B and recurrent parent DRR17B concerning other agro-morphological characters or grain type and the lines mostly resembled the original recurrent parent. The approach of coupling of MABB with phenotypic selection adopted in this study helped to regain most of the key agro-morphological traits of DRR17B, while simultaneously helping in the selection of some superior backcross derived segregants of DRR17B possessing BB resistance.

The improved lines of DRR17B were test crossed with IR58025A (WA-CMS line) to check their maintainer ability. Three lines (*viz*., IIRRIC102-26-7, IIRRIC123-34-84, and IIRRIC172-77-12) displayed complete maintainer ability highlighting the necessity of phenotypic confirmation for maintenance ability, despite a rigorous marker-assisted selection for *rf3* and *rf4* alleles in this study. This could be attributed to the existence of minor fertility restorer genes/QTLs as reported earlier [43].The three improved lines of DRR17B, possessing *Xa21 + Xa33* are being converted as CMS lines by crossing with DRR17A through MABB.

## Conclusion

The present study has resulted in the improvement of an elite maintainer of rice, DRR17B for durable resistance against BB, by incorporating two major dominant genes, *Xa21* and *Xa33* through marker-assisted backcross breeding (MABB). The two-gene pyramided lines of DRR17B displayed high levels of resistance against eight *Xoo* virulent isolates, comparable with three-gene pyramiding variety, Improved Samba Mahsuri (*Xa21*+*xa13*+*xa5*) and significantly better than the single gene containing lines (*Xa21* or *Xa33*). Three promising two-gene pyramiding lines of DRR17B with high level of BB resistance, agro-morphological attributes similar to or superior to the DRR17B with complete maintainer ability have been identified, and these lines are being converted to WA-CMS lines.

## Acknowledgment

The authors acknowledge Department of Biotechnology (DBT), Government of India, and Green Super Rice Project ICAR-Indian Institute of Rice Research, Hyderabad. The authors also thank the Project Director, ICAR-Indian Institute of Rice Research, for the provision of all the necessary facilities.

## Author contributions

SR planned and executed the experiment with the help of LG, HP, BS, SM, VB and GA. BC, BS and AV executed the experiment with the help of HG, HK, HS, DK, MA, KR, YA, PK, KM and RG. SK, VB and FA breeding team were involved in the selections. HP, SP and VB Hybrid rice breeding team were involved in the selections and conversion of selected improved lines into A line. LG, PM, and SR were involved in BB screening and scoring. BC, SR, LG, BM and AJ drafted the manuscript and conducted the statistical analysis. The manuscript was critically reviewed by SR, SM and AJ.

## References

1. Hariprasad AS, Viraktamath BC, and Mohapatra T. “Hybrid rice development in Asia: Assessment of limitations and potential”, In proceedings of regional expert conclusion (Bangkok). 2014; 85–100.

2. Indian Institute of Rice Research (IIRR). Production Oriented Survey (POS), Annual Progress Reports, All India Coordinated Rice Improvement Projects (AICRIP), Hyderabad 500030, India. 2008-2014.

3. Khush GS, and Jena KK. Current status and future prospects for research on blast resistance in rice (*Oryza sativa L.*). In: Advances in genetics, genomics and control of rice blast disease. 2009; 1–10. doi: https://doi.org/10.1007/978-1-4020-9500-9_1.

4. Mew TW. (1987). Current status and future prospects of research on bacterial blight of rice. Annu. Rev. Phytopathol. 1987; 25: 359–382.

5. Srinivasan B, and Gnanamanickam S. Identification of a new source of resistance in wild rice, *Oryza rufipogon* to bacterial blight of rice caused by Indian strains of *Xanthomonas oryzae* pv. *oryzae*. Curr. Sci. 2005; 88: 1229–1231.

6. Devadath S. Chemical control of bacterial blight of rice. In: Bacterial blight of rice. International Rice Research Institute, Manila, Philippines. 1989; 89–98.

7. Gnanamanickam S, Brindha PV, Narayanan N, Vasudevan P, and Kavitha S. An overview of bacterial blight disease of rice and strategies for its management. Curr. Sci. 1999; 77: 1435–1443.

8. Ellur RK, Khanna A, Yadav A, Sandeep P, Singh VK, Gopalakrishnan S,et al. Improvement of Basmati rice varieties for resistance to blast and bacterial blight diseases using marker-assisted backcross breeding. J. Plant Sci. 2015; 242: 330–341. doi: https://doi.org/10.1016/j.plantsci2015:08-020.

9. Sundaram RM, Chatterjee S, Oliva R, Laha GS, Leach JE, Cruz CV,et al. Update on bacterial blight of rice: Fourth International Conference on Bacterial Blight. Rice. 2014; 7: 12. doi: 10.1186/s12284-014-0012-7.

10. Kim SM, Suh JP, Qin Y, Noh TH, Reinke RF, and Jena KK. Identification and fine mapping of a new resistance gene, *Xa40*, conferring resistance to bacterial blight races in rice (*Oryza sativa* L.). Theor. Appl. Genet. 2015; 128: 1933–1943. doi: 10.1007/s00122-015-2557-2.

11. Hutin M, Sabot F, Ghesquiere A, Koebnik R, and Szurek B. A knowledge-based molecular screen uncovers a broad-spectrum *OsSWEET14* resistance allele to bacterial blight from wild rice. Plant J. 2015; 84: 694–703. doi: 10.1111/tpj.13042.

12. Laha GS, Reddy CS, Krishnaveni D, Sundaram RM, Srinivas PM, Ram T, et al. Bacterial Blight of Rice and Its Management. In: DRR Technical Bulletin No. 41. Directorate of Rice Research (ICAR), Hyderabad. 2009; p. 37

13. Shanti LM, Kumar VM, Premalatha P, Devi GL, Zher U, and Freeman W. Understanding the bacterial blight pathogen-combining pathotyping and molecular marker studies. Int. J. Plant. Pathol. 2010; 1: 58–68. doi: 10.3923/ijpp.2010.58.68.

14. Sundaram RM, Vishnupriya MR, Biradar SK, Laha GS, Reddy AG, Rani NS,et al. Marker assisted introgression of bacterial blight resistance in Samba Mahsuri, an elite *Indica* rice variety. Euphytica. 2008; 160: 411–422. doi: 10.1007/s10681-007-9564-6.

15. Hajira SK, Yugander A, Balachiranjeevi CH, Pranathi K, Anila M, Mahadevaswamy HK,et al. Development of durable bacterial blight resistant lines of Samba Mahsuri possessing *Xa33, Xa21, Xa13* & *Xa5*. Progressive Res. 2014; 9: 1224–1227.

16. Ronald PC, Albano B, Tabien R, Abenes MLP, Wu K, McCouch SR,et al. Genetic and physical analysis of the rice bacterial blight disease resistance locus *Xa21*. Mol. Gen. Genet. 1992; 236: 113–120.

17. Kumar PN, Sujatha K, Laha GS, Srinivasarao K, Mishra B, Viraktamath BC,et al. Identification and fine-mapping of *Xa33*, a novel gene for resistance to *Xanthomonas oryzae* pv. *oryzae*. Phyto Path. 2012; 102: 222–228. doi: 10.1094/PHYTO-03-11-0075.

18. Balachiranjeevi CH, Bhaskar NS, Abhilash V, Akanksha S, Viraktamath BC, Madhav MS,et al. Marker-assisted introgression of bacterial blight and blast resistance into DRR17B, an elite, fine-grain type maintainer line of rice. Mol. Breed. 2015; 35: 15. doi: 10.1007/s11032-015-0348-8.

19. Balaji SP, Srikanth B, Hemanth KV, Subhakara Rao I, Vemireddy L, Dharika N,et al. Fine mapping of *Rf3* and *Rf4* fertility restorer loci of WA-CMS of rice (*Oryza sativa* L.) and validation of the developed marker system for identification of restorer line. Euphytica. 2012; 187: 421–435. doi: 10.1007/s10681-012-0737-6.

20. Kauffman HE, Reddy APK, Hsieh SPY, and Merca SD. An improved technique for evaluating resistance of rice varieties to *Xanthomonas oryzae*. *Plant Dis*. Rep. 1973; 56: 537–540.

21. Lore JS, Vikal Y, Mandeep SH, Ravinder KG, Tajinder SB, and Girdhari LR. Genotypic and pathotypic diversity of *Xanthomonas oryzae* pv. *oryzae*, the cause of bacterial blight of rice in Punjab State of India. J. Pathol. 2011; 159: 479–487. doi: https://doi.org/10.1111/j.1439-0434.2011.01789.x.

22. IRRI. Standard Evaluation System for Rice (SES), 5th edition. Los Baños (Philippines): International Rice Research Institute. 2014.

23. Gomez KA, and Gomez AA. Statistical Procedures for Agricultural Research. New York, NY: John Wiley and Sons. 1984.

24. Ray DK, Mueller ND, West PC. and Foley JA. Yield trends are insufficient to double global crop production by 2050. PLoS One. 2013; 8: e66428. doi: https://doi.org/10.1371/journal.pone.0066428.

25. Ali J, Xu JL, Ismail AM, Fu BY, Vijaykumar CHM, Gao YM,et al. Hidden diversity for abiotic and biotic stress tolerances in the primary gene pool of rice revealed by a large backcross breeding program. Field Crops Res. 2006; 97: 66–76. doi: 10.1016/j.fcr.2005.08.016.

26. Basavaraj SH, Singh VK, Singh A, Singh A, Anand D, Yadav S,et al. Marker-assisted improvement of bacterial blight resistance in parental lines of Pusa RH10, a superfine grain aromatic rice hybrid. Mol. Breed. 2010; 26: 293–305. doi: 10.1007/s11032-010-9407-3.

27. Hari Y, Srinivasarao K, Viraktamath BC, Hariprasad AS, Laha GS, Ilyas A,et al. Marker-assisted improvement of a stable restorer line, KMR-3R and its derived hybrid KRH2 for bacterial blight resistance and grain quality. J Plant Breed. 2011; 130: 608–616. 608–616. doi: 10.1111/j.1439-0523.2011.01881.x.

28. Sundaram RM, Priya MRV, Laha GS, Rani NS, Srinivasarao, P, Balachandran SM,et al. Introduction of bacterial blight resistance into Triguna, a high yielding, mid-early duration rice variety by molecular marker assisted breeding. Biotechnol. J. 2009; 4: 400–407. doi: 10.10 02/biot.200800310.

29. Singh A, Singh VK, Singh SP, Pandian RTP, Ellur RK, Singh D,et al. Molecular breeding for the development of multiple disease resistance in Basmati rice. AoB Plant. 2012; pls029. doi: 10.1093/aobpla/pls029.

30. Hari Y, Srinivasarao K, Viraktamath BC, Hariprasad AS, Laha GS, Ahmed M,et al. Marker-assisted introgression of bacterial blight and blast resistance into IR 58025B, an elite maintainer line of rice. J. Plant Breed. 2013; 132: 586–594. doi: 10.1111/pbr.12056.

31. Khanna A, Sharma V, Ellur RK, Shikari AB, Gopalakrishnan S, Singh UD,et al. Development and evaluation of near-isogenic lines for major blast resistance gene(s) in Basmati rice. Theor. Appl. Genet. 2015; 128: 1243–1259. doi: 10.1007/s00122-015-2502-4.

32. Abhilash KV, Balachiranjeevi CH, Bhaskar NS, Rambabu R, Rekha G, Harika G,et al. Development of gene-pyramid lines of the elite restorer line, RPHR-1005 possessing durable bacterial blight and blast resistance. Front. Plant Sci. 2016; 7: 1195. doi: 10.3389/fpls.2016.01195.

33. Gizachew HG and Kumaravadivel N. Marker-assisted introgression of broad spectrum bacterial blight resistance gene *Xa33* into CO43, salt and alkaline soil tolerant Indica rice variety. Trends in Biosci. 2015; 8: 2136–2142.

34. Gizachew HG, Kumaravadivel N, Rabindran R, Ramanathan A, Soundararajan RP, and Selvi B. Identification and validation of microsatellite marker linked to the putative bacterial blight resistance gene *Xa33* in rice. Trends in Biosci. 2015; 8: 1069–1073.

35. Pradhan SK, Nayak DK, Mohanty S, Behera L, Barik SR, Pandit E,et al. Pyramiding of three bacterial blight resistance genes for broad-spectrum resistance in deep water rice variety, Jalmagna. Rice. 2015; 8: 19. doi: 10.1186/s12284-015-0051-8.

36. Bhaskar NS, Balachiranjeevi CH, Abhilash V, Harika G, Laha GS, Prasad MS,et al. Introgression of bacterial blight and blast resistance into the elite rice variety, Akshayadhan through marker-assisted backcross breeding. International J. Curr. Res. 2015; 7: 18943–18946.

37. Khush GS, Mackill DJ, and Sidhu GS. Breeding rice for resistance to bacterial blight. Bacterial Blight of Rice. Proceedings of the International Workshop on Bacterial Blight Rice, IRRI, Manila, Philippines. 1989; 207–217.

38. Shanti LM, George MLC, Cruz VCM, Bernando M, Nelson RJ, Leung H,et al. Identification of resistance genes effective against rice bacterial leaf blight pathogen. Plant Dis. 2001; 85: 506–512.

39. Mew TW, Cruz VCM, and Medalla ES. Changes in race frequency of *Xanthomonas oryzae* pv.*oryzae* in response to rice cultivars planted in the Philippines. Plant Dis. 1992; 76: 1029–1032.

40. Shanti LM, and Shenoy VV. Evaluation of BB resistance genes and their pyramids against rice bacterial leaf blight pathogen *Xanthomonas oryzae* pv*. oryzae*. Oryza. 2005; 42: 169–193.

41. Nayak D, Shanthi LM, Bose LK, Singh UD, and Nayak P. Pathogenicity association in *Xanthomonas oryzae* pv*. oryzae* the causal organism of rice bacterial blight disease. ARPN J. Agric. Biol. Sci. 2008; 3: 12–26.

42. Joseph M, Gopalakrishnan S, and Sharma RK. Combining bacterial blight resistance and Basmati quality characteristics by phenotypic and molecular marker-assisted selection in rice. Mol. Breed. 2004; 13: 377–387. doi: https://doi.org/10.1023/B:MOLB.0000034093.63593.4c.

43. Zhuang JY, Fan YY, Wu JL, Xia YW and Zheng KL. Mapping major and minor QTL for rice CMS-WA fertility restoration. Rice Genetics Newsletter. Research Notes-III. Genetics of physiological traits and others. 2000: 17: 56–58,

